# *Plap-1/Aspn* lineage tracing and single-cell transcriptomics reveals cellular dynamics in the periodontal ligament

**DOI:** 10.1101/2022.07.16.500314

**Authors:** Tomoaki Iwayama, Mizuho Iwashita, Kazuya Miyashita, Hiromi Sakashita, Shuji Matsumoto, Kiwako Tomita, Phan Bhongsatiern, Tomomi Kitayama, Kentaro Ikegami, Takashi Shimbo, Katsuto Tamai, Masanori A Murayama, Shuhei Ogawa, Yoichiro Iwakura, Satoru Yamada, Lorin E Olson, Masahide Takedachi, Shinya Murakami

## Abstract

Periodontal tissue supports teeth in the alveolar bone socket via fibrous attachment of the periodontal ligament (PDL). The PDL contains periodontal fibroblasts and stem/progenitor cells, collectively known as PDL cells (PDLCs), on top of osteoblasts and cementoblasts on the surface of alveolar bone and cementum, respectively. However, the characteristics and lineage hierarchy of each cell type remain poorly defined. This study identified *periodontal ligament associated Protein-1* (*Plap-1*/*Aspn*) as a PDL-specific extracellular matrix. We generated knock-in mice expressing CreER^T2^ and GFP specifically in *Plap-1*-positive PDLCs. Genetic lineage tracing confirmed the long-standing hypothesis that PDLCs differentiated into osteoblasts and cementoblasts. A PDL single-cell atlas defined cementoblasts and osteoblasts as *Plap-1*^-^*Ibsp*^+^*Sparcl1*^+^ and *Plap-1*^-^*Ibsp*^+^*Col11a2*^+^, respectively. Other populations such as *Nes*^*+*^ mural cells, S100B^*+*^ Schwann cells, and other non-stromal cells were also identified. RNA velocity analysis suggested that *Plap-1*^high^*Ly6a*^+^ cell population was the source of PDLCs. Lineage tracing of *Plap-1*^+^ PDLCs during the periodontal injury model showed periodontal tissue regeneration by PDLCs. Our study defines diverse cell populations in PDL and clarifies the role of PDLCs in periodontal tissue homeostasis and repair.

## Introduction

Mammalian teeth are supported by periodontal tissue, which consists of the periodontal ligament (PDL), gingiva, alveolar bone, and cementum. As periodontal tissue is highly adaptive to normal and excess occlusal and orthodontic forces, it undergoes extensive remodeling (Beertsen et al. 1997). Periodontal tissue also possesses some regenerative capacity, as demonstrated in human and animal models (Sallum et al. 2019; Kitamura et al. 2016; Nagayasu-Tanaka et al. 2015). The plasticity of periodontal tissue is thought to be underpinned by the mesenchymal stem/progenitor population in the PDL (Zhao et al. 2020). This population presumably gives rise to three stromal lineages: periodontal fibroblasts, cementoblasts, and osteoblasts (Luan et al. 2009; Iwayama et al. 2022). The PDL has also gained attention as a potent source of mesenchymal stem/stromal cells (MSCs) (Sharpe 2016) or multipotent stromal cells (Soliman et al. 2021). The PDL contains various other cell types, including epithelial cell rests of Malassez (Xiong et al. 2013), hematopoietic cells (Wilson et al. 2018), and neurovascular cells (Men et al. 2020). However, evidence regarding the lineage characteristics and cellular hierarchies of the adult mammalian PDL *in vivo* remains limited.

A reason for this ambiguity is the lack of clear phenotypic markers that distinguish the PDL fibroblast lineage (periodontal ligament cells (PDLCs), including stem/progenitor cells and fibroblasts) from other lineages of the periodontium. Based on recent lineage tracing analyses, many mesenchymal stem/progenitor markers have been proposed, including GLI-Kruppel family member GLI1 (Gli1)-, alpha-smooth muscle actin (αSMA)-, and Axin2-positive cells in the adult PDL (Roguljic et al. 2013; Yuan et al. 2018; Men et al. 2020). However, none of these markers are exclusively expressed in the PDL. Rather, they are also expressed in other components of the periodontium, as these markers are readouts of essential signaling pathways (Sonic hedgehog, transforming growth factor-β, and Wnt signaling, respectively) with dynamic expression in various tissues. Another challenge in PDL biology is the lack of efficient methods for isolating cells from tissues. Due to strong fiber attachments, dissociating the adult PDL is difficult (Jong et al. 2017). Recent single-cell transcriptome analyses of the periodontium have revealed that isolated cells predominantly comprise epithelial cells (Caetano et al. 2021) and constitute only a small number of PDL-derived stromal cells (Krivanek et al. 2020; Pagella et al. 2021). In this regard, a method for isolation and evaluation is required to decipher the cellular hierarchy of the PDL.

The PDL expresses a large number of extracellular matrix (ECM) components, including collagen (Naveh et al. 2018) and proteoglycans (Chen et al. 2021). Among these, several genes are candidate PDL-specific genes (Takimoto et al. 2015; Horiuchi et al. 1999). We previously identified *periodontal ligament associated Protein-1* (*Plap-1*/*Aspn*) from a human cDNA library and demonstrated that it is highly expressed in the PDL at the transcript and protein levels (Yamada et al. 2001; Yamada et al. 2007). Plap-1 is a 43-kDa extracellular matrix protein that is a subtype of the small leucine-rich family of proteoglycans (SLRP) (Kalamajski and Oldberg 2010). Plap-1 functions as a negative regulator of cytodifferentiation and mineralization of PDLCs through the interactions with BMP-2 and FGF-2 (Yamada 2006, Awata 2015). Its transcript levels in the maxilla, including the PDL, have been reported to be more than 10-fold greater than levels in adipose tissue, brain, heart, lung, liver, stomach, intestine, kidney, spleen, bone marrow, and muscle (Sakashita et al. 2021). Compared to dissected gingiva, dissected PDLs contain 60 times more transcripts (Ueda et al. 2021). Nevertheless, the expression of *Plap-1* in PDL at the single-cell resolution has yet to be investigated.

In this study, we identified *Plap-*1 as a molecule that clearly distinguished the PDL fibroblast lineage from other lineages, including osteoblasts and cementoblasts. Using knock-in mice expressing GFP and CreER^T2^ under the endogenous transcriptional control of *Plap-1*, we demonstrated that PDL cells differentiated into osteoblasts and cementoblasts *in vivo*. Further, we developed a method for highly efficient isolation of PDL-derived cells based on GFP expression in knock-in mice. A single-cell transcriptome atlas of the PDL revealed its cellular heterogeneity including *Plap-1*^+^*Nes*^+^ mural cells, and Schwann cells, and other non-stromal cells. Platelet-derived growth factor receptor alpha (PDGFRα)^+^ stromal cells in PDL could be divided into *Plap-1*^+^ PDLCs and *Bone sialoprotein* (*Ibsp)*^+^ cemento-/osteoblastic cells. Among *Ibsp*^+^ cells, *SPARC-like 1* (*Sparcl1)* and *Collagen type XI alpha 2* (*Col11a2)* were exclusively expressed in cementoblasts and osteoblasts, respectively. RNA velocity analysis suggested that *Plap-1*^high^Sca-1^+^ cells were the source of adult PDLCs. Upon tissue injury, PDLCs expanded to repair the periodontal tissue. These results advance our understanding of the PDL and its mechanisms of regeneration in both healthy and diseased states.

## Results

### Identification of *Plap-1* as a PDLC specific marker

The PDL is the extracellular matrix-rich connective tissue between the cementum and alveolar bone. Spindle-shaped fibroblastic cells were aligned along the horizontal axis of the fiber bundles (Fig. 1A). To identify exact cell types according to gene expression profiles at the single-cell resolution, we performed fluorescent *in situ* hybridization (FISH). We confirmed that both lineages expressed ECM components, such as *Collagen type I alpha 1* (*Col1a1), Biglycan (Bgn)*, and *secreted protein acidic and rich in cysteine* (*Sparc)* (Fig. 1B, 1C, and S1A). Cementoblasts and osteoblasts were labeled with *Secreted phosphoprotein 1* (*Spp1)* and *Ibsp* (Fig. 1D and 1F) as previously reported (Foster et al. 2018). We then found that only fibroblast-like cells in the PDL in the horizontal direction were positive for *Plap-1*, while the layer of cells on the surface of the alveolar bone and cementum was negative for *Plap-1* (Fig. 1E). FISH using both *Plap-1* and *Ibsp* probes showed that these cells are mutually exclusive in PDL (Fig. 1F). As Plap-1 is a secreted protein, its protein distribution was widespread throughout the PDL and could be seen around osteoblasts and cementoblasts in addition to the Plap-1 producing PDLCs (Fig. 1E vs. Fig.S1B). Having identified *Plap-1* as a molecular marker that specifically labels the periodontal fibroblast lineage (including stem/progenitor cells), we defined the three previously-proposed zones of the PDL (Lee et al. 2015) using *Plap-1* and *Ibsp* (Fig. 1G).

**Figure 1.**
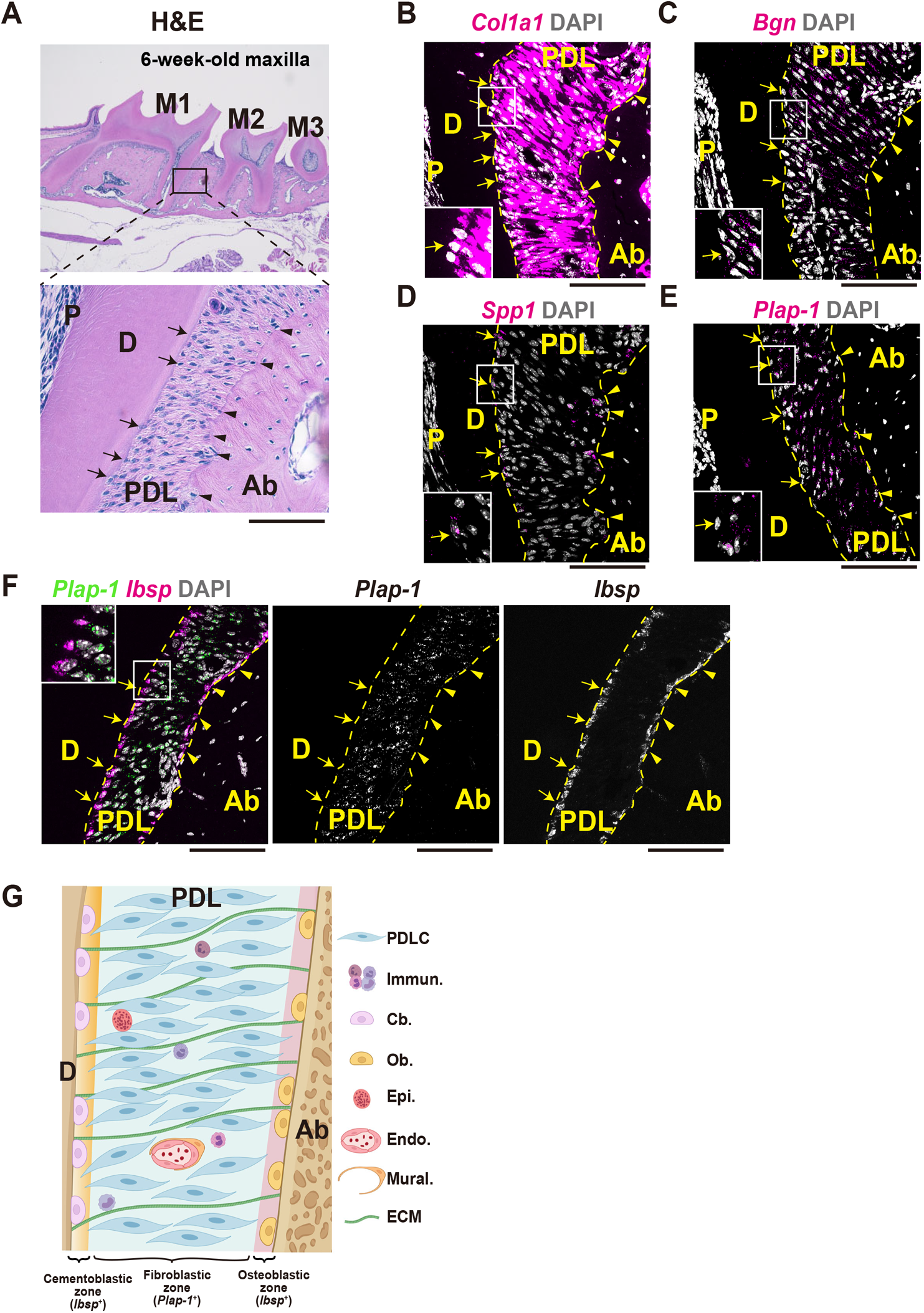
Fibroblastic and cemento-/osteoblastic zones in PDL A. Representative HE staining image of 6-week-old wild-type maxilla (lower magnification in upper panel) and PDL (higher magnification in lower panel). Arrows indicate cementoblasts. Arrowheads indicate osteoblasts. All PDL images in this paper are presented in this direction (left: cementum/dentin/root, right: alveolar bone). M1: maxillary first molar, M2: maxillary second molar, M3: maxillary third molar. B. Representative *Col1a1* mRNA expression in the PDL. Most cells in the PDL, including cemento-/osteoblasts, expressed *Col1a1*. C. Representative *Bgn* mRNA expression in the PDL. Similar with *Col1a1*, most cells in the PDL, including cemento-/osteoblasts, expressed *Bgn*. D. Representative *Spp1* mRNA expression in the PDL. *Spp1* -1 was expressed only in cells on the surface of the alveolar bone and cementum. E. Representative *Plap-1* mRNA expression in the PDL. In contrast to *Spp1, Plap-1* was expressed in most cells in the PDL but not in a subset of cells on the surface of the alveolar bone and cementum. F. Representative *Plap-1* (green) and *Ibsp* (magenta) mRNA expression in the PDL. The expression of *Plap-1* and *Ibsp* was mutually exclusive. G. Schematic representation of cellular composition in three zones of the PDL. The part in the cementum-alveolar bone direction is designated as the *Plap-1*^+^ fibroblastic zone. Either end (surface of cementum or alveolar bone) is designated as *Ibsp*^+^ cementoblastic or osteoblastic zone. PDLCs: PDL cells. Hem.: Hematopoietic cells. Epi.: Epithelial rests of Malassez. Endo.: Endothelial cells. Peri.: Pericytes surrounding endothelial cells. ECM: extracellular matrix. D: Dentin, Ab: Alveolar bone, P: Pulp, scale bars: 100 μm.

### Generation of PDLC-specific knock-in mice and lineage tracing

Next, we devised a strategy for CreER-2A-GFP knockin to faithfully reproduce *Plap-1* expression in mice (Fig. S1C-F). GFP expression in *Plap-1-GFP-2A-CreER* mice was consistent with the FISH results, and only cells in the fibroblastic zones were labeled (Fig. 2A). Furthermore, using the tamoxifen-induced Cre expression system, genetic lineage tracing of *Plap-1*^+^ cells can be conducted with the knockin mice (Fig. 2B). The short term lineage tracing revealed that they were restricted to the fibroblastic zone two days after treatment with tamoxifen (Fig. 2C) as expected. The tight regulation of CreER is further confirmed by no tdTomato expression without tamoxifen treatment (Fig. S1G) and limited tdTomato expression by a 1/10 dose tamoxifen treatment (Fig. S1H). The PDLCs were further traced for three months, when fibroblastic turnover of the tissue was completed (Sodek 1977). PDL still consisted of tdTomato^+^ cells, suggesting that the PDL homeostasis was maintained by *Plap-1*^+^ stem/progenitor cells (Fig. 2D). TdTomato^+^ cells were also observed both in cemento-/osteoblastic zones at three months post-treatment (Fig. 2D), suggesting that *Plap-1*^+^ cells gave rise to cemento-/osteoblast lineage cells. This was also confirmed by the existence of *Ibsp* expressing PDLC lineage cells in the middle and furcation areas of PDL (Fig. 2E). In addition, we investigated the Plap-1 expressions in the other tissues using the knockin mice (Fig. S2A-L), and confirmed the specificity of Plap-1 expression in PDL as previously reported (Sakashita et al. 2021).

**Figure 2.**
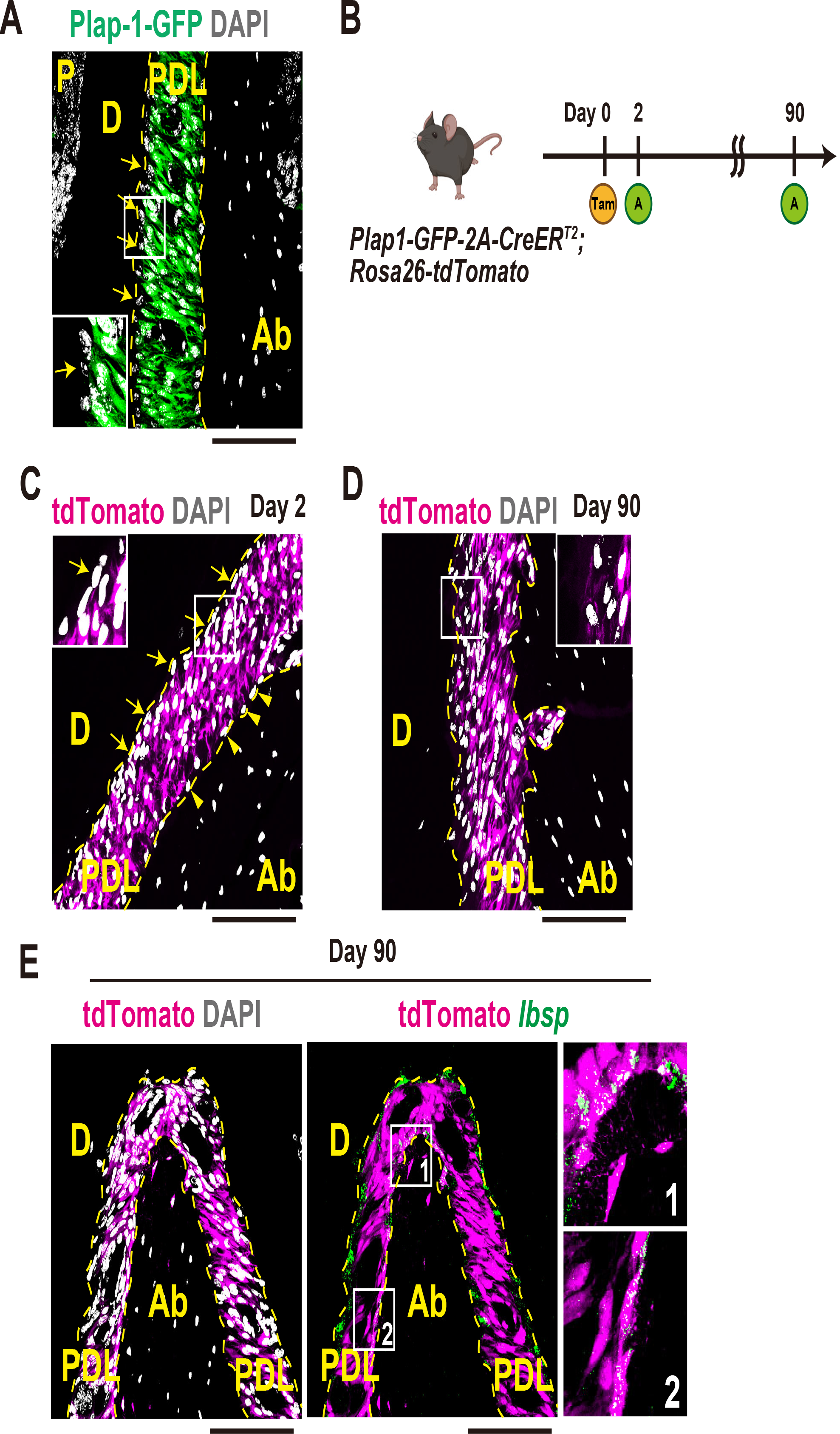
*Plap-1*^+^ cells maintain periodontal tissue A. Representative GFP expression in the PDL of *Plap-1-GFP-2A-CreER* knock-in mice. GFP expression was restricted to the fibroblastic zone. B. Overview of tamoxifen-induced lineage tracing experiment. *Plap-1-GFP-2A-CreER; Rosa26-tdTomato* mice were generated, and tamoxifen was administered. In the system, a fusion protein of a mutated estrogen receptor and Cre recombinase is expressed in Plap-1^+^ cells, and the Plap-1^+^ cells are labeled with tdTomato once tamoxifen is administrated. The mark is heritable and permanent and thus transmitted to all the descendants of Plap-1^+^ cells. T: Single tamoxifen administration via intraperitoneal injection. A: Analysis of the mice. C. Representative tdTomato expression in the PDL 2 days after tamoxifen treatment. *Plap-1* lineage cells were restricted to the fibroblastic zone. D. Representative tdTomato expression in the PDL 90 days after tamoxifen treatment. Cells in cemento-/osteoblastic zones were also positive for tdTomato. E. Representative tdTomato (magenta) and *Ibsp* mRNA (green) expression in the PDL of the furcation area 90 days after tamoxifen treatment. Cells in cemento-/osteoblastic zones and some osteocytes were also double-positive for tdTomato and *Ibsp*. Insets show *Ibsp* expressing Plap-1-lineage cells. D: Dentin, Ab: Alveolar bone, P: Pulp, scale bars: 100 μm.

### Development of a protocol for isolation of PDL-derived cells

To decipher the cell types in the PDL that contributes to the homeostasis and repair of periodontal tissue, an efficient method for isolation and analysis of PDL cells is required. We previously established a method for isolating PDL-derived cells by scraping the root surface with a PDL explant (Ueda et al. 2021). However, this method can only be used to analyze migratory cells. In this study, we used GFP expression in *Plap-1-GFP-2A-CreER* mice to evaluate the efficacy of PDL cell isolation. We adopted our previous protocol in which the gingiva was removed from the maxilla before tooth extraction to avoid any contamination from the gingival cells to extracted molars (Ueda et al. 2021). We first digested the scraped PDL with enzymes. Nevertheless, we did not observe any clear cell populations in the FSC/SSC plot (Fig. S3A). We then modified the protocol to digest extracted whole teeth (Fig. 3A) and observed a clear population in the FSC/SSC plot (Fig. S3A). Using flow cytometry and microscopy, we compared two enzymes, Liberase TM and Collagenase II. We identified a more evident population in the FSC/SSC plot with the use of Liberase TM (Fig. S3B). This was in agreement with the observation that the PDL was completely removed from the root surface with the use of Liberase TM (Fig. S3C). The histological analysis of the alveolar socket after tooth extraction showed that PDL tissue remained on the alveolar bone surface, but cells in the furcation area close to the alveolar crest were mostly removed (Fig S3D). We further confirmed that Liberase TM treatment was superior in terms of the percentage and the absolute number of live cells (Fig. S3E). Efficient PDL detachment from the tooth surface with Liberase TM was confirmed histologically (Fig. 3B). The isolated cells were analyzed using flow cytometry, and a gating strategy was established according to GFP expression.

**Figure 3.**
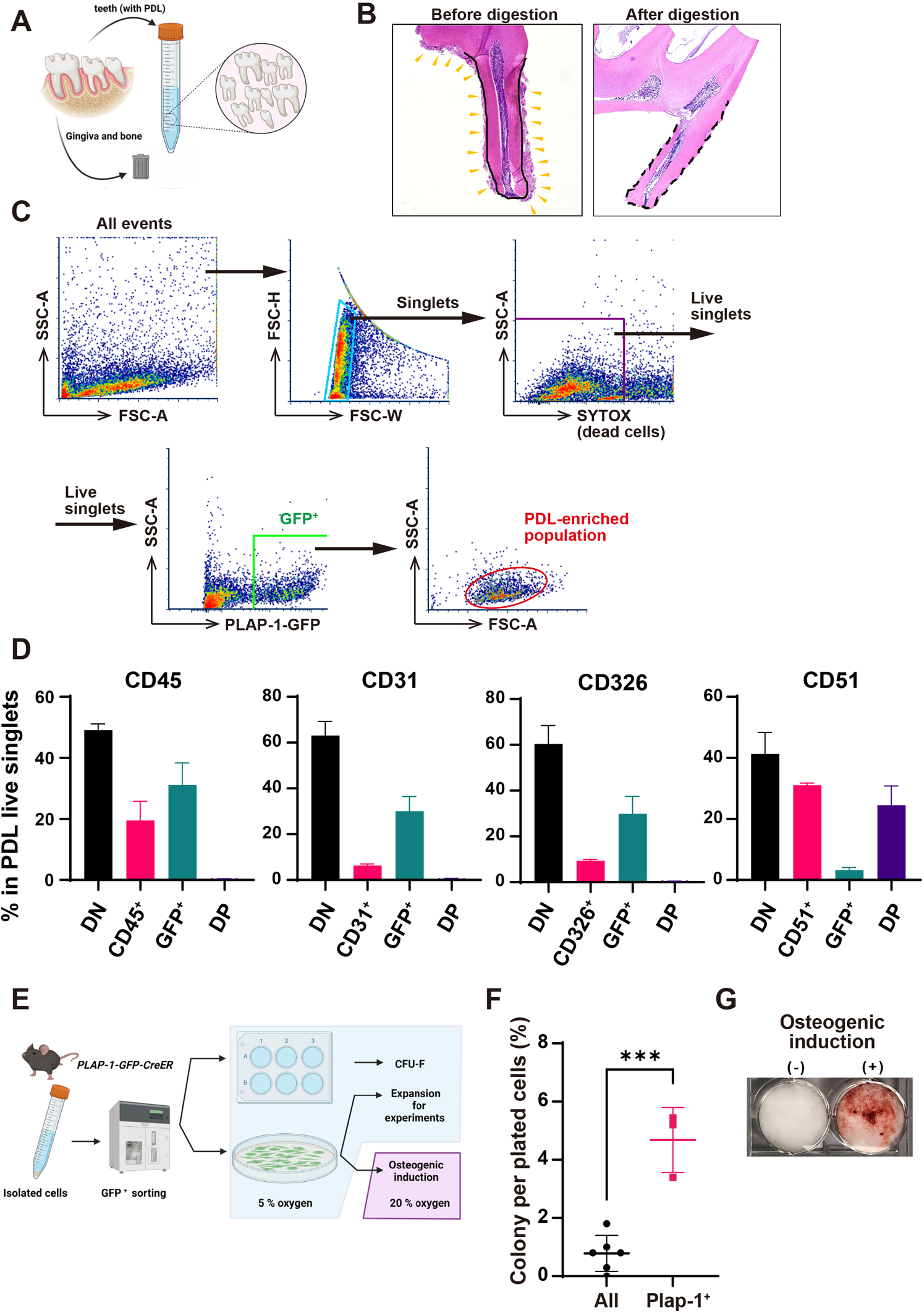
Efficient isolation and culture of PDL cells A. Overview of cell isolation experiments. After cardiac perfusion with PBS, the maxillae and mandibles were dissected, and gingival tissue was carefully removed under a stereomicroscope. The teeth were extracted from the remaining maxillae and mandibles and subjected to further processing. B. Representative HE staining of extracted teeth before and after Liberase TM digestion. The yellow arrowheads indicate PDL tissue in the left panel. In contrast, PDL tissue is absent but intact pulpal tissue is evident in the right panel. C. A gating strategy for PDLC-enriched populations by flow cytometry. Several populations were observed in FSC-A/SSC-A plots. Within live singlets, GFP^+^ PDLC was enriched in the green-colored gate shown in the FSC-A/SSC-A plot. D. Cellular composition analysis by flow cytometry. Within the PDL-enriched population, CD45, CD31, CD326, and CD51-positive cells were quantified (n=3 mice). DP: Double positive, DN: double negative. E. Overview of cell culture experiments. After isolation of PDL-derived cells from wild-type or *Plap-1-GFP-2A-CreER* mice, single-cell suspensions were subjected to cell sorting using an SH800 cell sorter. Sorted cells were expanded or analyzed using a CFU-F assay under hypoxia, followed by osteogenic induction experiments in normoxic conditions. F. CFU-F quantification. Plap-1-GFP^+^ cells and PDL-enriched populations were compared. The difference was statistically significant (p < 0.001, two tailed unpaired t-test, n= 3-6 mice, mean +/- S.D.). G. Representative Alizarin Red S staining images. After 20 days of induction, calcified nodules were observed only in the induction group.

GFP^+^ cells from live singlets were backgated onto the FSC/SSC plot, and the gating was defined as a PDL-enriched population (Fig. 3C). The population was then used to determine the relative proportion of each cell type. PDL cell-enriched populations yielded more than 10,000 cells per animal, of which 20%, 6%, and 10% comprised CD45^+^ immune cells, CD31^+^ vascular endothelial cells, and CD326^+^ epithelial cells, respectively (Fig. 3D). In addition, we investigated whether a stromal cell marker CD51, also known as integrin αV, can label stromal cells including Plap-1^+^ cells. CD51^+^ cells included most GFP^+^ cells, with an average of 30% single-positive cells. GFP^+^ cells consistently accounted for approximately 30% of the population. We also tested the workflow using a Col1a1-GFP mouse line that expresses GFP in both fibroblastic and cemento-/osteoblastic cells in periodontal tissue (Fig. S3F). As expected, more GFP^+^ cells were observed in the Col1a1-GFP PDL than in the Plap-1-GFP PDL (Fig. S3G). A comparison of gingiva with the PDL revealed that Col1a1-GFP expression levels were higher in the PDL than in gingiva, consistent with a previous report that collagen turnover is more active in the PDL than in gingiva (Sodek 1977). Additionally, we examined the culture conditions suitable for PDLCs and developed a method that enabled stable PDL cell expansion *in vitro* under hypoxic conditions (Fig. 3E). GFP^+^ cells exhibited significantly higher CFU-F formation compared to all cells from the PDL-enriched population (Fig. 3F) and demonstrated osteogenic differentiation capacity *in vitro* (Fig. 3G), confirming that PDLCs contained stem/progenitor cells. However, cellular composition and heterogeneity of PDL were not determined from the analysis.

### Generation of PDL atlas using single-cell transcriptome analysis

After establishing an efficient PDLC isolation technique, we obtained cell suspensions from 20 adult wild-type mice. All viable cells in the PDL were collected by cell sorting and subjected to single-cell transcriptome analysis to generate a “PDL single-cell atlas” (Fig. 4A, 4B, and S4A). After the removal of low-quality cells, 17 clusters were identified from the PDL scRNA-seq data (7,318 cells) (Fig. 4C). The cell types in each cluster were annotated using known cell marker genes (Fig. 4D). The most abundant populations were stromal cells expressing collagens, matrix metalloproteinases, and platelet-derived growth factor receptors (Fig. 4D and S4B). Consistent with the histological analysis (Fig. 1F), collagen-expressing stromal cells were mutually exclusively defined by the expression of *Plap-1/Aspn* or *Ibsp* (Fig. 4E). The stromal cell cluster was subjected to the in-depth analysis (Fig. 5). Non-stromal cell clusters, including immune, epithelial, and vascular endothelial cells, were also identified (Fig. 4D, S4C, and S4E). Flow cytometry analysis of isolated immune cells from PDL showed the presence of macrophages, proerythroblasts, B cells, and T cells in PDL and no overlap with Plap-1-GFP^+^ cells (Fig. S4D). Two epithelial cell clusters were distinguished by differential expression of *Keratin 15* (*Krt15)* and *Keratin 17 (Krt17)*. FISH analysis confirmed that *Krt17-*positive epithelial cells were derived from the inner and outer epithelia, while *Krt15-*positive epithelial cells were derived from the basement membrane (Fig. S4F). Notably, a cluster of *Plap-1*^+^ cells exhibited distinct gene expression in stromal cell clusters. This cluster expressed *Nes, Regulator of G-protein signaling 5* (*Rgs5)*, and *Myosin-11* (*Myh11)* (Fig. 4E F), which are markers of vascular mural cells (vascular smooth muscle cells and pericytes), suggesting that the PDL also contains mural cell populations that are less heterogeneous than the fibroblast lineage (Muhl et al. 2020). To identify *Plap-1*^+^*Nes*^+^ cell clusters in PDL histology, we used *Nes-GFP* transgenic mice, in which stem/progenitor populations are labeled in other tissues (Méndez-Ferrer et al. 2010; Iwayama et al. 2015). Nes-GFP^+^ cells exhibited a perivascular appearance, with long cellular processes along the blood vessels in the PDL (Fig. 4G). The cell bodies were covered by the basement membrane of vascular endothelial cells, suggestive of pericytes (Fig. 4H).

**Figure 4.**
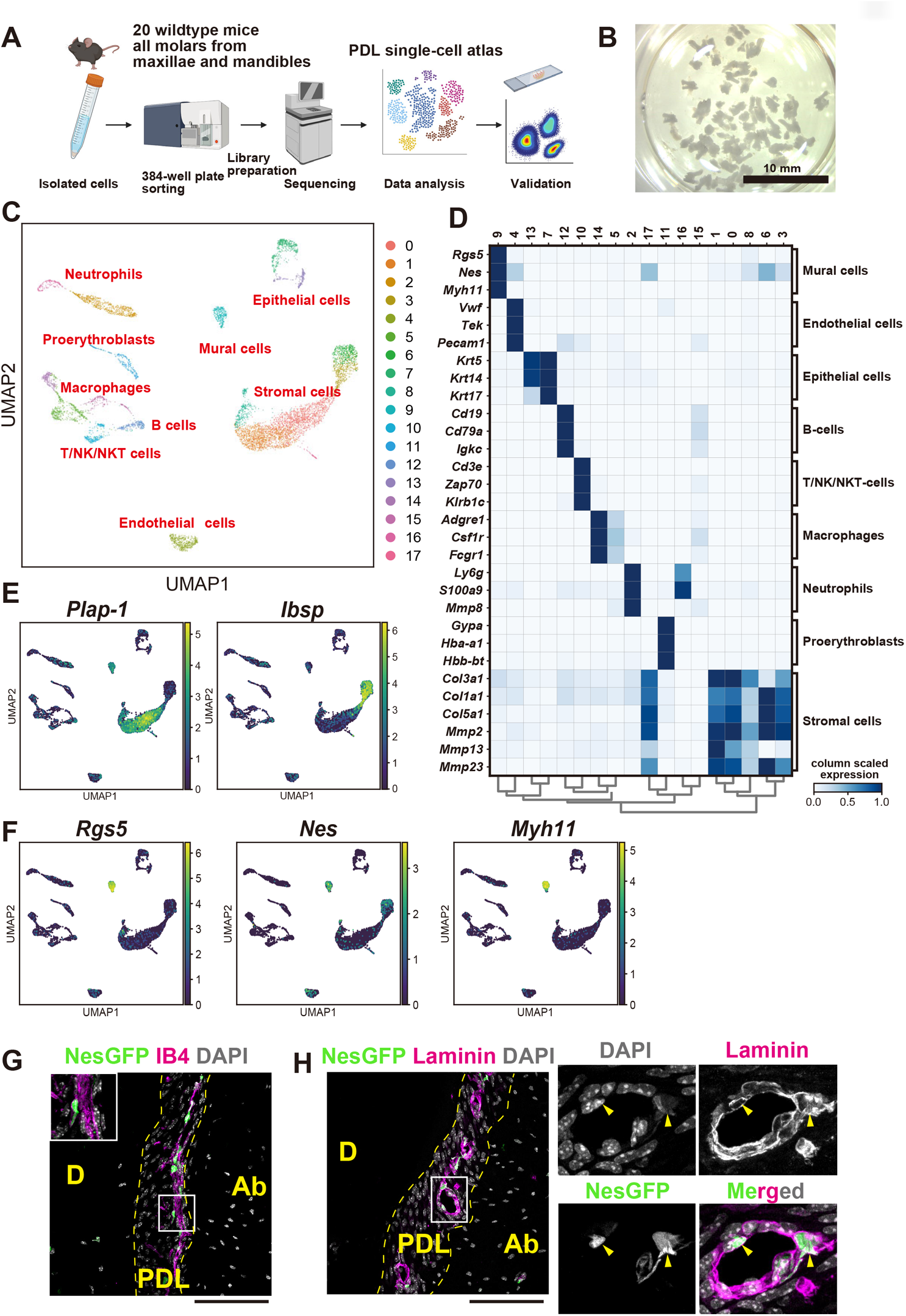
Single-cell transcriptomic analysis of PDL-derived cells A. Overview of the experimental workflow. Teeth from 20 wild-type mice were subjected to cell isolation, and each cell type was sorted into 384 well plates with a lysis solution. Each lysate was then processed for library preparation, followed by sequencing and bioinformatic analysis. The identified molecules were validated using flow cytometry or histological analysis. B. Representative photograph of extracted teeth used for scRNA-seq analysis. All molars were extracted from dissected maxillae and mandibles. C. scRNA-seq data obtained from 7,318 PDL-derived cells. Clustering analysis with 17 cell types is shown in UMAP. D. A heatmap of the expression of representative genes for each cluster. Each cell type was annotated using these marker genes. E. UMAP plots showing *Plap-1* and *Ibsp* expression. The expression of *Plap-1* and *Ibsp* was mutually exclusive. F. UMAP plots showing *Rgs5, Nes*, and *Myh11* expression. The expressions of *Rgs5, Nes*, and *Myh11* were restricted to mural cell cluster. G. Representative GFP expression and GS-IB-4 staining in the PDL of Nestin-GFP mice. Insets showed Nestin-GFP^+^ cells located in the perivascular area in the PDL. H.Representative laminin immunofluorescent staining in the PDL of Nestin-GFP mice. Nestin-GFP^+^ cell bodies were encapsulated by the endothelial basement membrane. D: Dentin, Ab: Alveolar bone, scale bars: 100 μm.

**Figure 5.**
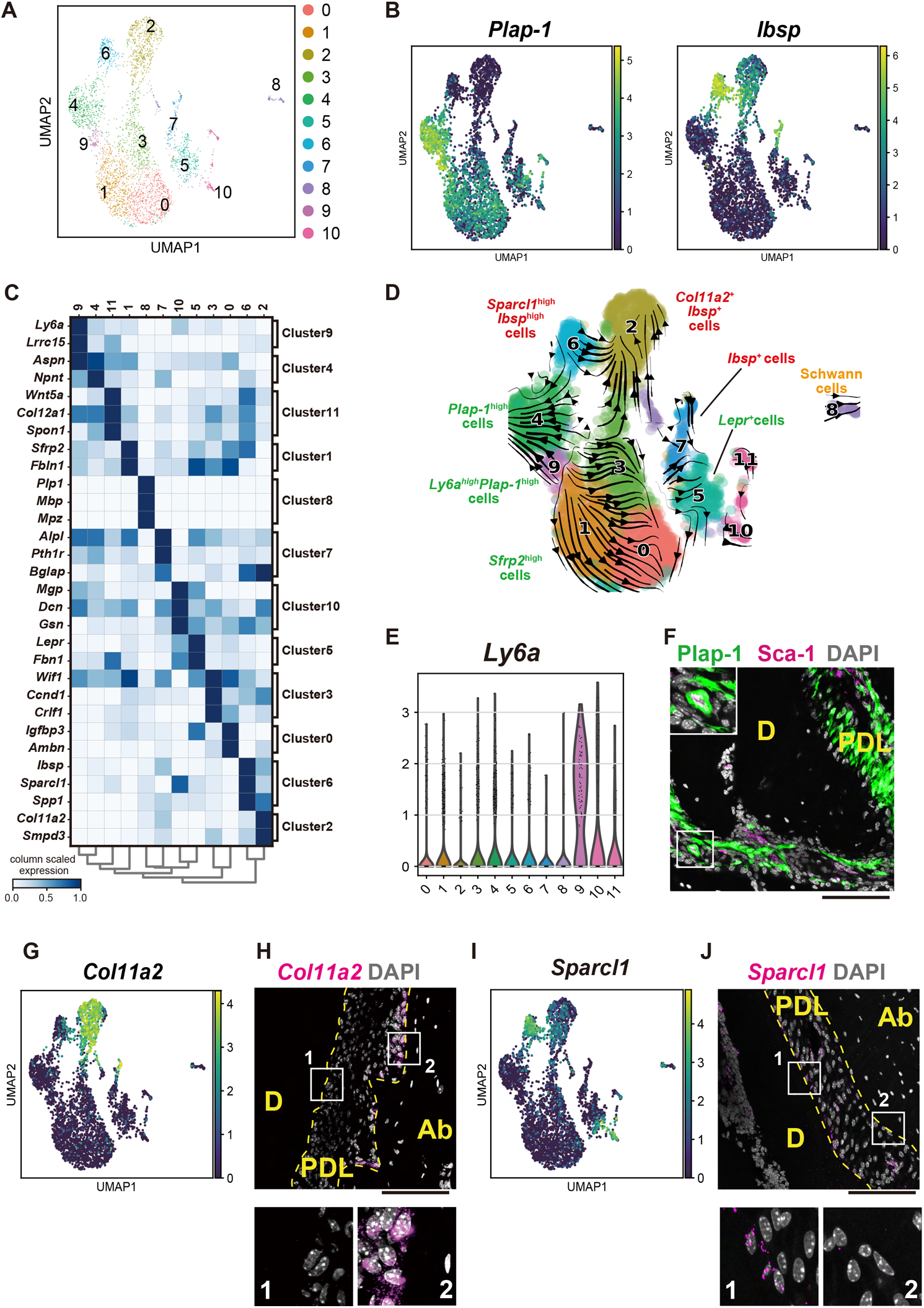
Sub-clustering analysis of PDL stromal cells A. Sub-clustering analysis of stromal cells. B. UMAP plots showing *Plap-1* and *Ibsp* expression. The expression of *Plap-1* and *Ibsp* was mutually exclusive. C. A heatmap for representative genes for each cluster. D. RNA velocity analysis of the sub-cluster. RNA velocity in every single cell was estimated by distinguishing between unspliced and spliced mRNAs, whose balance is predictive of cellular state progression. E. Violin plot showing Ly6a expression in each cluster. The expression was specifically high in cluster 9. F. Representative Sca-1 immunofluorescent staining in the PDL. Sca-1^+^ PDL cells were located in the apical area of the PDL (inset). *G. Col11a2* expression overlayed on UMAP plot. The expression was specifically high in cluster 2. H. Representative *Col11a2* mRNA expression in the PDL. The expression was restricted to the osteoblastic zone. H. *Sparcl1* expression overlayed on UMAP plot. The expression was specifically high in cluster 6. I. Representative *Sparcl1* mRNA expression in the PDL. The expression was restricted to the cementoblastic zone.

### Cellular hierarchy of the PDL according to RNA velocity analysis

To identify the heterogeneity in the PDL that maintains tissue homeostasis, 3,675 stromal cells expressing collagens, matrix metalloproteinases, and platelet-derived growth factor receptors (Fig. S4B) were subjected to subclustering analysis. We identified 11 clusters (Fig. 5A) and confirmed that they were expressed either *Plap-1* or *Ibsp*, except cluster 8 (Fig. 5B). Intriguingly, *Scx, Mkx*, and *Tnmd*, which have been previously reported to be PDLC-specific, were expressed only in a subset of both *Plap-1*^+^ and *Ibsp*^+^ cells (Fig. S5A). It should be noted that the master regulators of osteogenic differentiation, Runt-related transcription factor 2 (*Runx2)* and *Sp7 transcription factor 7 (Sp7*, as known as osterix), were also globally expressed in both lineages, suggesting that PDLCs are readily differentiated into osteoblasts and cementoblasts (Fig. S5B). This skewed differentiation is also supported by the expression of other bone-related proteins and SLRPs in PDLCs (Fig. 1B, 1C, and S1A). Based on the representative gene expression profile (Fig. 5C), cluster 8 was annotated as Schwann cells, and S100 calcium-binding protein B (S100B) expressing cells were identified in PDL (Fig. S5C). In addition to the annotation of each cell type within stromal cells, we conducted RNA velocity analysis that can predict the future state of individual cells by distinguishing between unspliced and spliced mRNA (Fig. 5D). The result suggested that the cells in cluster 9 as the source of differentiated cells in the PDL environment. This cluster showed higher expression and positive velocity of *Plap-1* (data now shown). In search of the marker genes, *lymphocyte activation protein-6a* (*Ly6a)* and *Leucine rich repeat containing 15* (*Lrrc15*) were identified. *Ly6a* encodes Stem cell antigen (Sca-1) that has been used for identifying stem cells in many tissues in combination with another antigen, such as PDGFRα (Houlihan et al. 2012). The relative expression level of the Ly6a is higher in the cluster (Fig. 5E), and we further characterized the population as Sca-1-positive *Plap-1*^high^ cells.

Histological analysis revealed that they were located in the apical part of the PDL (Fig. 5F), and flow cytometry analysis showed they were 5.4 % of the PDL live singlets (Fig. S5D). *Lepr-, Gli1*-, *Axin2*-, *and Acta2*-positive cells have been implicated in stem/progenitor cells in the PDL. However, in this dataset, they were not associated with a specific cluster, with the exception of *Lepr*^+^ cells that were enriched in cluster 5 (Fig. S5E). Crucially, none of these cells were observed at the top of the lineage hierarchy (Fig. 5D). Among *Ibsp*^+^ clusters, *Col11a2* expression was specific to cluster 2 (Fig. 5G), and this gene was exclusively expressed in osteoblasts but not in cementoblasts (Fig. 5H). Compared to the osteoblast cluster, the other *Ibsp*^+^ cluster (cluster 6) expressed higher levels of *Sparcl1* (Fig. 5I). *Sparcl1* expression was observed in cementoblasts and cementocytes, but not in osteoblasts (Fig. 5J and S5F). However, the cluster did not specifically express recently reported markers (Nagata et al. 2021), including *Parathyroid hormone-related protein* (*Pthlh), Class III β-tubulin* (*Tubb3)*, and *Wnt inhibitory factor 1 (Wif1)* (Fig. S5G). To assess the putative differentiation pathways, we also performed a pseudotime analysis to clusters 0, 2, 4, and 6. Gene expression kinetics of *Plap-1, Ibsp, Col3a1, Col1a1, Sparcl1, Col11a2, Lrrc15*, and *Ly6a* showed unique transient changes during differentiation (Figure S6A-F).

### *Plap-1*^+^ PDLCs directly contribute to periodontal tissue repair

To investigate the role of PDLCs in the repair of injured periodontal tissue, we utilized a ligature-induced periodontitis model (Fig. 6A). In this widely used model (Abe and Hajishengallis 2013), a suture is placed around the second molar of the maxilla for 7 days to induce periodontal tissue loss. Upon ligature removal, the periodontal tissue can be repaired to the baseline level 7-14 days after removal. On day 1, after ligature removal, a layer of cementum on the surface remained intact, although alveolar bone resorption was observed (Fig. 6B, left). On day 7, recovered alveolar bone levels and fibrous attachment between the bone and cementum were observed (Fig. 6B, right). *Plap-1* mRNA expression was restricted to the fibroblastic zone between the remaining alveolar bone and cementum (Fig. 6C), which was defined as the PDL. Notably, *Ibsp* expression was dramatically downregulated in the remaining cementum (Fig. 6D). To elucidate the contribution of *Plap-1*^+^ cells in remaining PDL to tissue repair, *Plap-1-GFP-2A-CreER; Rosa26-tdTomato* mice were administered tamoxifen 1 day before ligature removal (Fig. 6E). In the early stages of periodontal wound healing, the granulation tissue beneath the gingival epithelial layer consisted of *Plap-1* lineage cells and αSMA-positive myofibroblasts (Fig. 6F and G). The 20.2 +/- 6.7 % of αSMA-stained cells in granulation tissue was Plap-1 lineage cells (n=6, standard deviation), which was 22.4 +/- 6.7 % of entire Plap-1 lineage cells in the granulation tissue (n=6, standard deviation). These data suggest the partial contribution of the Plap-1^+^ cells to the granulation tissue formation. On day 7, in addition to most PDLCs in repaired PDL, 56.5 % of osteoblasts and cementoblasts on the surface of repaired alveolar bone and cementum were derived from Plap-1^+^ lineage cells (Fig. 6H-J). These data confirm the long-standing concept that periodontal tissue is maintained and repaired by the PDL and emphasize the importance of the PDL.

**Figure 6.**
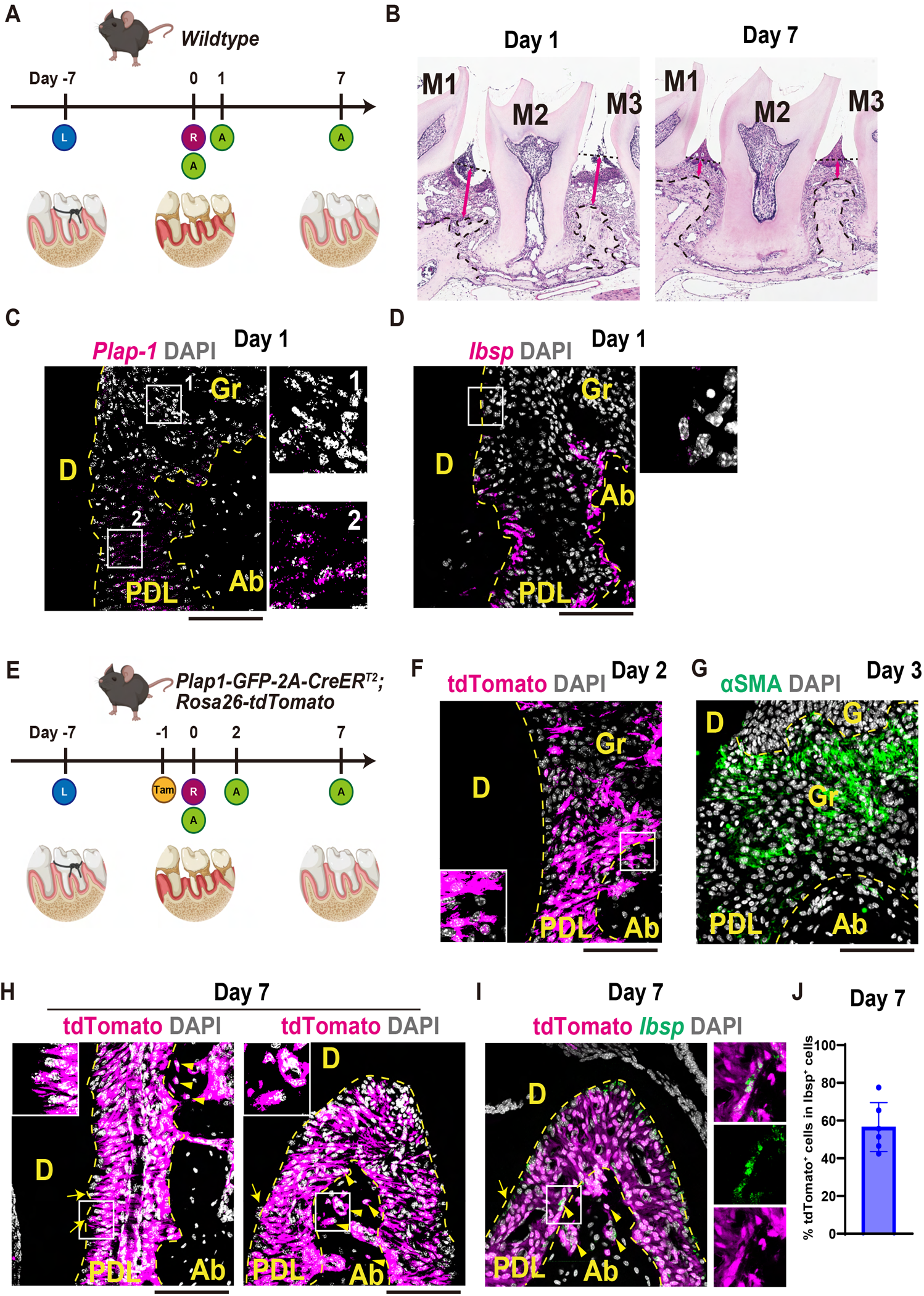
*Plap-1* lineage cells in repaired periodontal tissue A. Overview of ligature-induced periodontitis experiments. Eight-week-old wild-type mice underwent ligature placement around the second molar of the maxilla. After 7 days, periodontal tissue was inflamed and destroyed. A week after suture removal, periodontal tissue was repaired. L: Ligature placement. R: removal of suture. A: Analysis of the mice. B. Representative HE staining image of periodontal tissue destruction. On the day of suture removal (day 0), the periodontal tissue was destroyed. The tissue consisted of newly formed granulation tissue and remaining periodontal tissue. The arrow showed a distance between cemento-enamel junction and the alveolar bone crest. C. Representative *Plap-1* mRNA expression in the border of the remaining PDL and granulation tissue. Only the remaining PDL (defined by the presence of neighboring bone and cementum) expressed *Plap-1*. D. Representative *Ibsp* mRNA expression in the border of the remaining PDL and granulation tissue. *Ibsp*^+^ cementoblasts on the cementum surface were not evident beyond the border of the remaining PDL and granulation tissue. E. Overview of ligature-induced periodontitis experiments. *Plap-1-GFP-2A-CreER; Rosa26-tdTomato* mice were administered tamoxifen and underwent ligature placement around the second molar of the maxilla. The suture was removed 7 days after ligature placement (designated as day 0). The repair process was analyzed on days 2 and 7. A week after suture removal, periodontal tissue was repaired. L: Ligature placement. R: removal of suture. A: Analysis of the mice. F. Representative tdTomato expression in repairing periodontal tissue 2 days after suture removal. Plap-1-lineage cells were distributed in granulation tissue. G. Representative αSMA immunofluorescent staining in the granulation tissue 2 days after suture removal. αSMA^+^ cells were distributed in granulation tissue, but only 20.2 % were derived from Plap-1-lineage cells based on quantification within histological sections. H. Representative tdTomato expression in repairing periodontal tissue 7 days after suture removal. The repaired tissue consisted of *Plap-1* lineage cells. Of note, the osteocytes were differentiated from PDLCs (arrowheads). I. Representative tdTomato (magenta) and *Ibsp* mRNA (green) expression in the PDL of the furcation area 7 days after suture removal. Some cells were double-positive for tdTomato and *Ibsp* (highlighted with arrows for cementoblasts and arrowheads for osteoblasts). Insets show Ibsp expressing Plap-1-lineage cells. J. Quantification of tdTomato^+^ cells in Ibsp-expressing cells within histological sections. D: Dentin, Ab: Alveolar bone, Gr: Granulation tissue, scale bars: 100 μm.

### Mural cell population in the PDL is quiescent cells

Long-term lineage tracing analysis with Nes-Cre revealed that Nes^+^ mural cells were quiescent and remained perivascular for up to a year without expansion during periodontal tissue homeostasis (Fig. S7A). Based on this observation, we performed a ligature experiment with Nes-Cre lineage tracing to determine if *Plap-1*^+^*Nes*^+^ cells contribute to periodontal tissue repair (Fig. S7B). On day 7, after ligature removal, we did not detect the *Nes* lineage cells in repaired PDL, alveolar bone, and cementum (Fig. S7C). These results suggest that *Nes*^*+*^ mural cells in the PDL did not contribute to the periodontal tissue repair in a cell-autonomous manner.

## Discussion

Periodontitis is a chronic inflammation of periodontal tissues caused by bacterial biofilms, which eventually cause irreversible destruction of periodontal tissues. Periodontitis has a high prevalence worldwide and is the most common cause of tooth loss in adults in developed countries. In addition to attaching teeth to the alveolar bone, the PDL interferes with and senses the occlusal force and transmits it to the central nervous system (Beertsen et al. 1997). Dental implants, however, lack these functions because they form a direct connection to the alveolar bone (osseointegration) without the PDL. Fibrous attachment of the PDL is one of the most clinically important indicators of periodontal disease and regeneration. It has long been proposed that the PDL contains stem/progenitor populations based on the fact that periodontal tissue can partially regenerate (Kitamura et al. 2016; Sallum et al. 2019) and has a large adaptive capacity for orthodontic treatment and occlusal forces in addition to remodeling capacity (Beertsen et al. 1997). PDL tissue attached to extracted human teeth has been analyzed extensively, and PDL stem cells (PDLSCs) with minimal stemness criteria (Mao et al. 2012), including *in vitro* differentiation capacity to osteoblasts and ectopic bone formation by transplantation assays, have been widely used (Li et al. 2021). However, as with MSCs in other tissues, the definition of PDLSCs is ambiguous, and rigorous *in vivo* analyses are required to define these cell types. In this study, we established a PDL single-cell atlas and demonstrated that the PDL comprises heterogeneous cell types. Notably, the stromal compartment was collectively divided into *Plap-1*^+^ fibroblastic cells or *Ibsp*^+^ osteo-/cementoblastic cells in a mutually exclusive manner. Plap-1 is also highly expressed in the human PDL and used as a cultured PDLC marker (Yamada et al. 2001; Loo-Kirana et al. 2021; Mun et al. 2022). RNA velocity analysis indicated that cluster 9, *Plap-1*^high^, with a positive velocity of *Plap-1* mRNA, constituted a potential cellular source of other PDL clusters. This population expressed *Ly6a* which encodes Sca-1, which is known to be expressed in the stem cell population and in epithelial and endothelial cells. This *Plap-1*^high^ *Ly6a*^+^ population could be PDLSCs that maintain and repair the entire periodontal tissue and overlap with the recently defined skeletal stem cell-like population in periodontal tissue (Liang et al. 2022). To clarify the cell fate of the putative PDLSC *in vivo*, a definitive single marker gene or an intersectional lineage tracing approach will be needed in future studies.

In this study, we also harnessed single-cell analysis in combination with Nes-GFP and Nes-Cre transgenic mice to demonstrate the presence of strictly defined mural cells in the PDL for the first time. Although the association of pericytes with MSCs in other tissues has been described (Soliman et al. 2021), our findings suggest that *Nes*^+^ cells are quiescent mural cells near capillaries, and they did not directly contribute to PDL regeneration. Further study is needed to clarify if they have an indirect contribution, including the release of paracrine signals and exosomes (Wagoner and Zhao 2021). Our single-cell analysis revealed the presence of mesenchymal cells and many immune cells, especially macrophages and monocytes, in the PDL. The presence of CD45-positive immune cells has been reported (Soliman et al. 2021), but this remains inconclusive due to the lack of efficient methods to isolate PDLCs and other cells from the PDL. Future studies should elucidate how pericytes and PDLCs repair tissues by interacting with immune cells at each stage of wound healing. Other cell types, such as plasma cells that were not analyzed in this study (Fig. S4C), might be identified by specific isolation of immune cells and deeper sequencing.

Regeneration of the cementum-PDL-alveolar bone complex is a major goal of periodontal therapy. In contrast to bone regeneration, the mechanism of cementum regeneration remains unclear. Without regeneration of the cementum, new attachment formation by the PDL cannot be obtained even if the bone has regenerated. In the ligature model, *Ibsp*^+^ cementoblasts disappeared from the root surface of destroyed periodontal tissue. *In vivo* lineage tracing of PDLCs demonstrated that *Ibsp*^+^ cells reappeared in the regenerated tissue and more than half of *Ibsp*^*+*^ cells originated from the PDLCs (Fig. 6H-J). However, the exact cellular dynamics of the cementoblasts were to be investigated with definitive markers. Recent analyses of cementum development in mice have highlighted several molecules that are specifically expressed in cementoblasts (Nagata et al. 2021). In this study, we defined cementoblasts as *Plap-1*^-^*Ibsp*^+^*Sparcl1*^+^ and alveolar osteoblasts as *Plap-1*^-^*Ibsp*^+^*Col11a1*^+^. Sparcl1 is an ECM protein related to Sparc and is secreted by nerves and blood vessels (Naschberger et al. 2016; Fan et al. 2021). In the future, single-cell epigenetic profiling of periodontal tissue may be harnessed to identify transcriptional regulators of cementoblasts.

In conclusion, our work identified *Plap-1* as a PDLC-specific molecule, and cell lineage analysis using knock-in mice confirmed that PDLCs differentiated into osteoblasts and cementoblasts and played a role in periodontal tissue homeostasis. In addition, we developed a novel cell isolation method and performed scRNA-seq to generate a PDL single-cell atlas that clearly defined osteoblasts and cementoblasts. Furthermore, lineage tracing revealed that injured periodontal tissues were repaired by PDLCs. These findings will contribute to the development of efficient regenerative therapy for periodontitis.

## Acknowledgments

We thank Ms. Kotomi Aso and Ms. Minako Mori for her technical assistance. The Nes-GFP mouse line was a gift from Dr. Grigori Enikolopov of Stony Brook University. The Col1a1-GFP mouse line was a gift from Dr. David Brenner of the University of California, San Diego. Confocal microscopy was performed at the Center for Frontier Oral Science (Osaka University Graduate School of Dentistry). Flow cytometry was performed at the Center for Medical Research and Education (Graduate School of Medicine, Osaka University). Figures were created using Biorender.com. This study was supported by the Japan Society for the Promotion of Science (JSPS) KAKENHI Grant Numbers JP15J03981, JP16H06273, JP17K19752, JP19H03830, JP19K22713, and JP19H01069.

## Author contributions

Conceived and designed the experiments: TI and SM. Performed the experiments: TI, MI, HS, SM, KT, PB, TK, MAM, SO, YI, SY, LEO, and MT. Analyzed the data: TI, KM, KI, TS, and KT. Drafted the manuscript: TI, MI, KM, TK, and TS. Critically revised the manuscript: MAM, SO, YI, SY, LEO, MT, and SM.

## Lead contact

Further information and requests for resources and reagents should be directed and will be fulfilled by the lead contact, Tomoaki Iwayama

## Materials availability

Unique materials generated in this study are available from the lead contact without restriction.

## Data and code availability

- Single-cell RNA sequencing data were accessed from GEO: GSE197828.
- All data needed to evaluate the conclusions in the paper are presented in the paper and/or Supplementary Information.
- Additional data related to this study can be requested from the authors.

## Experimental model and subject details

### Mice

All animal experiments in this study were approved by the Animal Experimentation Committee of Osaka University Graduate School of Dentistry (approval numbers: R-02-012-0 and 31-001-0). *Plap-1-GFP-2A-CreER* mice were generated using a conventional gene-targeting strategy. *R26-LSL-tdTomato* (stock #007909) and *Nes-Cre* (stock #003771) were obtained from Jackson Laboratory (Bar Harbor, ME, USA). *Nes-GFP* mice were provided by Dr. Grigori Enikolopov of Stony Brook University. *Col1a1-GFP* mouse were provided by Dr. David Brenner, University of California, San Diego. PCR genotyping was performed using the primers listed in Supplementary Table S1. Mice were provided sterile food and water under specific-pathogen-free conditions. All animal experiments complied with the ARRIVE guidelines.

### Method details

#### ES cell targeting and vector design

A targeting vector harboring the PGK-DTA-long homology arm-GFP-2A-CreER-WPRE-pA-short homology arm sequence was generated. Correctly targeted ES cell clones were identified by PCR and confirmed by Southern blotting using the primers and probes listed in Supplementary Table S1. Chimeric males were generated using standard aggregation of clumped ES cells with eight-cell embryos and transplantation procedures and were used to establish germline transmission.

#### Preparation of paraffin and frozen sections

Mice were euthanized with CO_2_ gas, perfused, and fixed with 4% paraformaldehyde/phosphate buffer (PFA; Fujifilm Wako Pure Chemicals, Osaka, Japan, #163-20145). Maxillary bones and other tissues were harvested, and after overnight immersion fixation in PFA, the hard tissue was demineralized in demineralizing solution B (Fujifilm Wako Pure Chemicals, #041-22031) for 3-5 days with three exchanges of solution. For paraffin sections, the tissue was embedded in paraffin and sectioned at 7 μm.

HE-staining was performed as previously described (Ueda et al. 2021). For frozen sections, the tissue was immersed in 20% sucrose (Fujifilm Wako Pure Chemicals, #196-00015) solution and then embedded in TissueTech O.C.T. compound (Sakura Finetech Japan, Tokyo, Japan, #4583), frozen in ethanol cooled on dry ice. Tissue sectioning at 14 μm was performed using a cryostat CM3050S (Leica Microsystems, Wetzlar, Germany). A cryofilm (type 2C(9), Section-lab, Hiroshima, Japan) was used as section support.

#### Fluorescent immunohistochemistry

The prepared frozen sections were washed with PBS and blocked with Blocking One Histo (Nacalai Tesque, Kyoto, Japan, #06349-64) for 30 min at room temperature. Sections were incubated with rabbit anti-mouse laminin polyclonal primary antibody (1:1,000, Sigma-Aldrich, St Louis, MO, USA, #L9393) at 4°C overnight followed by incubation with Alexa Fluor 647-conjugated goat anti-rabbit IgG secondary antibody (1:200, Thermo Fisher Scientific, Waltham, MA, USA, #A32733) at room temperature for 30 min. For myofibroblast identification, sections were incubated with AlexaFluor 647 conjugated anti-mouse αSMA monoclonal antibody (1:100, clone 1A4, Santa Cruz Biotechnology, Dallas, TX, USA, #sc-32251) at 4°C overnight. Nuclear and vascular staining was performed with DAPI (1:50,000, Sigma-Aldrich, #D9542) and Alexa Fluor 647-conjugated Isolectin GS-IB4 (1:200, Thermo Fisher Scientific, #I32450), respectively. Fluorescently labeled tissue sections were imaged using a fluorescence microscope (ECLIPSE Ti, Nikon, Tokyo, Japan) or Leica TCS SP8 confocal microscope (Leica Microsystems, Wetzlar, Germany). For confocal microscopy, 10-14 z-stack images (total thickness of approximately 5 µm) were acquired, and 3D images were obtained using the maximum value projection method.

#### RNAscope fluorescent in situ hybridization

Cryosections were hydrated and hybridized (Advance Cell Diagnostics, Newark, CA, USA) using the Multiplex Fluorescent Reagent v2 kit according to the manufacturer’s instructions. Nuclei were counterstained with DAPI (1:50,000, Sigma-Aldrich, #D9542), sealed with ProLong Diamond Antifade Mountant (Thermo Fisher Scientific, #P36961), and observed under a Leica TCS SP8 confocal microscope, as described above. The probes used in this study are listed in Supplementary Table S1.

### Tamoxifen treatment

Tamoxifen (Sigma-Aldrich, #T5648) was dissolved in corn oil at a concentration of 20 mg/mL and mixed overnight. Mice (6-8 weeks old) received an intraperitoneal injection of 100 μL of tamoxifen (2 mg). In low-dose experiments, 10 µL of tamoxifen (0.2 mg) was administered intraperitoneally and analyzed after 16 days. 14-week-old Plap-1-GFP-2A-CreER; Rosa26-tdTomato mice without tamoxifen treatment were used to check any leaky expression of tdTomato.

### Ligature-induced periodontitis model

To induce experimental periodontitis, a 5-0 silk ligature was tied around the left maxillary second molar of 8-week-old mice as described previously (Ueda et al. 2021). After 7 days, the ligature was removed. Tissue recovery was analyzed on day 7 after removal. The contralateral molar tooth in each mouse was left unligated as intact periodontal tissue.

### Isolation of PDL-derived cells

After euthanization with CO_2_ gas, the mice were perfused with PBS, and the maxilla and mandible were separated. Gingival tissue was peeled and carefully removed. Subsequently, the molars were extracted and collected into PBS in a 12-well plate. These procedures were performed under a stereomicroscope. The teeth were washed three times and transferred to fresh wells. Liberase TM (Merck, Darmstadt, Germany, #05401119001) or collagenase type II (Worthington Biochemical, Lakewood, NJ, USA, #CLS-2) solution was used to digest PDL, as described in the Results section. Liberase TM powder was reconstituted with 2 mL of distilled water and stored at -20°C. The working solution was prepared in digestion buffer (PBS containing 2% FCS) at a final concentration of 2 Wünsch Units/mL. The teeth were then incubated in 2 mL digestion buffer in 15 mL low-binding tubes (STEMFULL; Sumitomo Bakelite, Tokyo, Japan, #MS-90150) for 20 min at 37°C and 200 rpm. Subsequently, 20 µL of 500 mM EDTA was added, and the teeth were further incubated for 10 min at 37°C and 200 rpm. After digestion, 10 mL of buffer was added, and cells were centrifuged at 400× *g* for 8 min at 4°C. The pellet was resuspended in 2 mL of buffer and passed through a 70-µm cell strainer (BMS, Tokyo, Japan, #BC-AMS-17001). After centrifugation at 320× *g* for 5 min at 4°C, cells were finally resuspended in 400 µL of buffer and subjected to downstream analyses. In some experiments, PDL was scraped from the surface of the mesial root of the first molar using micro-instruments, including a micro-curette (0.5 mm) (Fine Science Tools). Scraped tissue was digested as described above.

### Flow cytometry analysis

The isolated cells were resuspended in FACS buffer (PBS containing 5% FBS). The cell pellet was stained for 30 min on ice with fluorescent protein-conjugated antibodies (Supplementary Table S1). Flow cytometry analysis was performed on a FACSCanto II (BD) flow cytometer using FCS Express 7 (De Novo Software) or FlowJo v10.8 Software (BD).

### Sorting and primary culture of PDL-derived cells

Cell sorting was performed using an SH800Z Cell Sorter (Sony Biotechnology, CA, USA) with 100-μm microfluidic sorting chips. The isolated cells were expanded and maintained at 37ºC, 5% O_2_, and 10% CO_2_ with a MesenCult Expansion Kit (STEMCELL Technologies, ST-05513) according to the manufacturer’s protocol. The culture medium contained MesenPure, L-glutamine (Wako, #073-05391), and penicillin-streptomycin (Wako, #168-23191). For the CFU-F assay, 500-1000 sorted cells were maintained in a 6-well plate. On day 14, the cells were fixed with methanol and stained with Giemsa staining solution (Nacalai Tesque, #37114-64). Fibroblastic colonies with more than 20 cells were quantified as CFU-F.

### Osteogenic differentiation culture

To induce osteogenic differentiation, confluent cells were cultured in osteogenic medium comprising cell culture medium supplemented with 50 µg/mL l-ascorbic acid phosphate magnesium salt n-hydrate (Fujifilm Wako Pure Chemical, #013-19641) and 10 mM glycerol 2-phosphate disodium salt n-hydrate (Fujifilm Wako Pure Chemical, #193-02041). The medium was replaced with fresh medium every 3 days. For mineral staining, cells were fixed with cold 100% ethanol for 5 min and stained with 1.0% Alizarin Red S solution (pH adjusted to 6.4; Wako Pure Chemical Industries, Japan).

### scRNA-seq

Single-cell RNA-seq was performed based on a previous report with the following modifications. Primer mix composed of 5 μL of lysis buffer comprising 3.1375 μL of Buffer EB (QIAGEN, #19086), 0.5 μL of 10 mM dNTP (GenScript, Piscataway, NJ, USA, #C01582), 0.05 μL of Phusion HF buffer (New England BioLabs (NEB), Beverly, MA, #B0518S), 0.3125 μL of Proteinase K (Nacalai Tesque, # 15679-64), and 1 μL of 1 μM barcoded oligo-dT primer (5’-ACGACGCTCTTCCGATCT[Barcode]NNNNNNNNTTTTTTTTTTTTTTTTTTTTTTTTTTT TTTVN-3’, where “N” is any base and “V” is either “A”, “C” or “G”; IDT) was aliquoted into 384-well plates. The isolated PDL-derived cells were stained with SYTOX™ Blue dead cell stain (Thermo Fisher Scientific, #S34857). Unstained cells were considered live cells. Live cells were sorted into plates using a BD FACSAria III instrument (BD Biosciences; 100-µm chip) in single-cell purity mode. The plates were immediately centrifuged and frozen at -80°C. Plates were incubated at 50°C for 10 min and then at 80°C for 10 min. A volume of 5 μL of first-strand reaction mix containing 2 μL of 5× Superscript IV First-Strand Buffer (Thermo Fisher Scientific, #18090050B), 0.5 μL of 100 mM DTT (Thermo Fisher Scientific), 0.0125 μL of SuperScript IV reverse transcriptase (200 U/μL, Thermo Fisher Scientific, #18090200), 0.05 μL of SUPERase In RNase Inhibitor (Thermo Fisher Scientific, #AM2696), and 2.4375 μL of water was aliquoted into each well. The plates were then incubated at 55°C for 10 min, and the reaction was inactivated by incubation at 80°C for 10 min. To remove unincorporated oligos, 2 μL of exonuclease I mix containing 0.0625 μL of exonuclease I (Thermo Fisher Scientific, #EN0582), 1.2 μL of 10× reaction buffer, and 0.7375 μL of water was added, and the mixture was incubated at 37°C for 20 min. Samples were pooled and purified using a DNA Clean & Concentrator Kit-100 (Zymo Research, Orange, CA, USA, #D4030), concentrated using a DNA Clean & Concentrator Kit-5 (Zymo Research, #D4014), and eluted in 12 μL of Buffer EB. Eluted cDNA was denatured at 95°C for 2 min and immediately placed on ice for 2 min. The cDNA was pre-amplified using the Accel-NGS 1S Plus DNA Library Kit (#10096; Swift Biosciences, Ann Arbor, MI, USA). A volume of 10 μL of Adaptase reaction mix containing 1.25 μL of Buffer EB, 2 μL of Buffer G1, 2 μL of Reagent G2, 1.25 μL of Reagent G3, 0.5 of μL Enzyme G4, 0.5 μL of Enzyme G5, and 0.5 μL of Enzyme G6 was added, and the mixture was incubated at 37°C for 15 min and at 95°C for 2 min. A volume of 23.5 μL of Extension reaction mix containing 9.25 μL of Buffer EB, 1 μL of Reagent Y1, 3.5 μL of Reagent W2, 8.75 μL of Buffer W3, and 1 μL of Enzyme W4 was added, and the mixture was incubated at 98°C for 30 s, 63°C for 15 s, and 68°C for 5 min. Amplified cDNA was purified using 26.1 µL of AMPure XP beads (Beckman Coulter Diagnostics, Brea, CA, USA, #A63881) and eluted in 19.5 μL of Buffer EB. To amplify cDNA libraries, each well was mixed with 3 μL of 10 μM i5 primer (5’-AATGATACGGCGACCACCGAGATCTACAC[i5]ACACTCTTTCCCTACACGACGCTCTT CCGATCT-3’; IDT), 2.5 μL of 12 μM D7 primer (CAAGCAGAAGACGGCATACGAGATCGAGTAATGTGACTGGAGTTCAGACGTGTGCTC TTCCGATC-3’; IDT), and 25 μM KAPA HiFi ReadyMix (KAPA Biosystems, Boston, MA, USA, #KK2602). Amplification was performed using the following program: 98°C for 3 min; 14 cycles of 98°C for 20 s, 67°C for 15 s, and 72°C for 2 min; and a final hold at 72°C for 5 min. Each well was purified using 30 μL of AMPure XP beads, eluted in 30 μL of Buffer EB, and transferred to a new PCR tube. Each well was then purified using 18 μL of AMPure XP beads and eluted in 10 μL of buffer EB. Amplified cDNA (1 ng) was mixed with water to a total volume of 5 μL. The following reactions were performed using the Nextera XT DNA Library Prep Kit (Illumina, San Diego, CA, USA; #FC-131-1096). Each well was mixed with 10 μL of Nextera TD buffer and 5 μL of Amplicon Tagment enzyme and then incubated at 55°C for 5 min for tagmentation. After tagmentation, samples were mixed with 20 μL of DNA Binding Buffer #D4004 -1-L; Zymo Research), purified using 32 μL of AMPure XP beads, and eluted in 16 μL of Buffer EB. The eluted DNA was mixed with 2 μL of 10 μM i5 primer, 2 μL of 10 μM P7 primer (5’-CAAGCAGAAGACGGCATACGAGAT[i7]GTCTCGTGGGCTCGG-3’), and 20 μM NEBNext High-Fidelity 2× PCR Master Mix (NEB, #M0541L). Amplification was performed using the following program: 72°C for 3 min, 98°C for 30 s, 8 cycles of 98°C for 10 s, 66°C for 30 s, and 72°C for 1 min, and a final hold at 72°C for 5 min. After amplification, the samples were purified using 32 μL of AMPure XP beads and eluted in 10 μL of Buffer EB. Purified samples were quantified for concentration using the Qubit dsDNA HS Assay Kit (Thermo Fisher Scientific, #Q32854), and size was measured using a High Sensitivity D5000 Reagent Kit (Agilent Technologies, Santa Clara, CA, USA, #5067-5593) and High Sensitivity D5000 Screen Tape (Agilent Technologies, #5067-5592). Sequencing libraries were sequenced on a NextSeq2000 platform (Illumina). The read length was set to 20 (read 1), 8 (i7), 8 (i5), and 51 (read 2) bases.

### scRNA-seq data processing

BCL files were obtained from NextSeq (Illumina, San Diego, CA) and then demultiplexed and converted into FASTQ files using bcl2fastq2 (https://jp.support.illumina.com/downloads/bcl2fastq-conversion-software-v2-20.html).

FASTQ reads were mapped onto mouse mm10 reference genomes using STARv2.7.6a with solo options (Dobin et al. 2013). The count tables from STAR were loaded as a Seurat object (Hao et al. 2021). Low-quality samples with extremely high (>3500) or low (<200) UMI counts or high percentages of mitochondrial genes (>10%) were removed. Downstream analyses, including normalization, scaling, and clustering, were conducted using Seurat v4 functions following a standard analytic procedure. For sub-cluster analysis, stromal cells were extracted from whole cells, and normalization and scaling were performed. A phenograph was employed in the sub-cluster analysis to run more detail clustering (Levine et al. 2015). Gene expression dynamics along pseudotime were calculated using Palantir (Setty et al. 2019).

### RNA velocity

Spliced and unspliced transcripts were quantified with Velocyto (La Manno et al. 2018) using BAM files from STARv2.7.6, with *the --soloFeatures Gene GeneFull SJ Velocyto* option. The output loom files were combined with the Seurat object and then converted into an h5ad file using SeuratDisk (https://mojaveazure.github.io/seurat-disk/). Further RNA velocity analyses were performed using the Scanpy (Wolf et al. 2018) and scVelo (Bergen et al. 2020) packages.

### Quantification and statistical analysis

Data are presented as mean ± standard deviation (SD) in Fig. 3F, 6J and S6D or hmean ± standard error (SEM) in Figs. 3D, S3E and S4D otherwise raw data values were plotted. Statistical analysis was performed using GraphPad Prism v9.3 (GraphPad Software, USA, www.graphpad.com). Statistical significance was assumed at p < 0.05. Quantification of the tdTomato, αSMA, and Ibsp mRNA was performed by counting positive cells in the targeted area (day 3: granulation tissue upon ligature was defined as the tissue between gingival epithelium and repaired alveolar crest, day 7: PDL attached to coronal half of the root) of histological sections. One field of view each from two sections from three individual mice was used for the counting cells.

